# SARS-CoV-2 Serosurvey of healthy, privately owned cats presenting to a New York City animal hospital in the early phase of the COVID-19 pandemic (2020-2021)

**DOI:** 10.1101/2024.02.13.580068

**Authors:** Annette Choi, Alison E. Stout, Alicia Rollins, Kally Wang, Qinghua Guo, Javier A. Jaimes, Monica Kennedy, Bettina Wagner, Gary R. Whittaker

**Affiliations:** Departments of Microbiology & Immunology, Ithaca NY and Sutton Animal Hospital, New York NY; Public & Ecosystem Health, Ithaca NY and Sutton Animal Hospital, New York NY; Population Medicine & Diagnostic Sciences, Ithaca NY and Sutton Animal Hospital, New York NY; College of Veterinary Medicine, and Cornell Public Health Program, Ithaca NY and Sutton Animal Hospital, New York NY; Cornell University, Ithaca NY and Sutton Animal Hospital, New York NY

## Abstract

SARS-CoV-2, the cause of the ongoing COVID-19 pandemic, not only infects humans but is also known to infect various species, including domestic and wild animals. While many species have been identified as susceptible to SARS-CoV-2, there are limited studies on the prevalence of SARS-CoV-2 in animals. Both domestic and non-domestic cats are now established to be susceptible to infection by SARS-CoV-2. While serious disease in cats may occur in some instances, the majority of infections appear to be subclinical. Differing prevalence data for SARS-CoV-2 infection of cats have been reported, and are highly context-dependent. Here, we report a retrospective serological survey of cats presented to an animal practice in New York City, located in close proximity to a large medical center that treated the first wave of COVID-19 patients in the U.S. in the Spring of 2020. We sampled 79, mostly indoor, cats between June 2020 to May 2021, the early part of which time the community was under a strict public health “lock-down”. Using a highly sensitive and specific fluorescent bead-based multiplex assay, we found an overall prevalence of 13/79 (16%) serologically-positive animals for the study period; however, cats sampled in the Fall of 2020 had a confirmed positive prevalence of 44%. For SARS-CoV-2 seropositive cats, we performed viral neutralization test with live SARS-CoV-2 to additionally confirm presence of SARS-CoV-2 specific antibodies. Of the thirteen seropositive cats, 7/13 (54%) were also positive by virus neutralization, and two of seropositive cats had previously documented respiratory signs, with high neutralization titers of 1/1024 and 1/4096; overall however, there was no statistically significant association of SARS-CoV-2 seropositivity with respiratory signs, or with breed, sex or age of the animals. Follow up sampling of cats showed that positive serological titers were maintained over time. In comparison, we found an overall confirmed positive prevalence of 51% for feline coronavirus (FCoV), an endemic virus of cats, with 30% confirmed negative for FCoV. We demonstrate the impact of SARS-CoV-2 in a defined feline population during the first wave of SARS-CoV-2 infection of humans, and suggest that human-cat transmission was substantial in our study group. Our study provide a new context for SARS-CoV-2 transmission events across species.

**Significance:** SARS-CoV-2 has a broad animal tropism and can infect a wide range of animal species, leading to an expansion of the viral reservoir. Expansion of this viral reservoir may result in the accumulation of mutations within these species, potentially giving rise to new viral variants and facilitating reverse zoonotic transmission. Domestic cats are particularly noteworthy in this regard due to their close contact with humans. Currently, there are very limited studies on the prevalence of SARS-CoV-2 infection in domestic cats during the early stages of the pandemic, especially in the United States. This retrospective study addresses the gap by investigating seroprevalence of SARS-CoV-2 in cats in New York City, the epicenter of the COVID-19 pandemic in the United States during the early pandemic. Our work underscores the importance of adopting a One Health approach to pandemic prevention and conducting routine surveillance across different animal species

## Introduction

SARS-CoV-2, the causative agent of the COVID-19 pandemic, is a Betacoronavirus that has caused significant morbidity and mortality in the human population since its emergence in December 2019 (1,2). SARS-CoV-2 has relatively broad cellular tropism and utilizes angiotensin converting enzyme 2 (ACE2) as a primary receptor to enter host cells. ACE2 is highly conserved across many mammals—which, in part, may explain why SARS-CoV-2 is also able to infect a range of animal species.

Following a first report in dogs (3), it became apparent that additional species were becoming infected with SARS-CoV-2, including tigers and lions at the Bronx Zoo in New York City in early April 2020 (4). As the pandemic has proceeded most cases in cats have been found to be subclinical; the virus may circulate between cats (5,6), but the public health risk from cats is now considered low— and while transmission back to humans can occur, this is rare. Other examples of cross-species infection with SARS-CoV-2 have now been widely documented, with natural infections also reported in farm-raised American mink, otters, hamsters, and white-tailed deer, among other species (7–10).

Cats, in particular, are of special interest due to their close contact with humans and predicted receptor-based susceptibility to SARS-CoV-2. Feline ACE2 shares significant homology with its human ACE2, especially at the spike receptor binding interface, or RBD (11,12). Despite initial *in vitro* studies, the overall impact of spike RBD/ACE-2 interactions on susceptibility to infection may ultimately be quite low. Experimental studies have, however, demonstrated cats high susceptibility to SARS-CoV-2 infections and their potential to transmit the virus through close contact (13). Numerous studies have now reported seroprevalence of SARS-CoV-2 in domestic, shelter, and stray cats, yielding variable prevalence (14–29). Despite the now extensive global documentation, data on the seroprevalence of SARS-CoV-2 in U.S. domestic cats remains relatively scarce (25–29), especially during the early stages of the pandemic.

Detecting active infection in animal species can often be challenging, as in some species SARS-CoV-2 infection is asymptomatic. Serological assays are an effective method for detecting past SARS-CoV-2 infections in animals species and inferring infection history. Additionally, serological assays support a ‘passive surveillance’ approach, which minimizes bias by not restricting sample selection solely to those with known prior exposure. This allows for a comprehensive assessment of seroprevalence across all cats, offering a holistic understanding of SARS-CoV-2 prevalence in animal species.

Here, we present a retrospective serological survey conducted in a small animal veterinary clinic located on the Upper East Side of New York City, in close proximity to a large medical center (New York Presbyterian/Weill Cornell Medicine) that was treating the initial U.S. wave of severe COVID-19 cases in humans. Our sampling period spanned June 2020 to May 2021, a period that included an early epicenter of the COVID-19 pandemic in the U.S., with numerous human cases identified spanning the New York City boroughs by early March of 2020 (30,31). Given the dense population of both humans and pets in this area, our findings provide key insights into the prevalence and characteristics of SARS-CoV-2 infection in domestic cats during the initial outbreak phase.

## Materials and Methods

### Participant recruitment and sampling

Feline participants presenting to a private practitioner veterinarian in New York City were enrolled between June 2020 and May 2021. Serum samples from a total of 79 domestic cats (33 male, 46 female) were collected. The average age of the cats was 7.3 years. From each cat, approximately 2 mL of whole blood was collected and allowed to coagulate. Serum was removed, stored at 4 °C and shipped to Cornell University, Ithaca, NY on ice. Sampling from cats was approved by the Cornell IACUC 2012-0116, and owners gave their written consent for this procedure. All sampling was performed by licensed veterinary professionals, employed by the collaborating veterinary practice. Basic signalment was collected from each cat. Positive samples identified in this study were reported to New York State Department of Agriculture and Markets, Division of Animal Industry and United States Department of Agriculture, Veterinary Services, and were submitted to the National Veterinary Services Laboratory in Ames, Iowa for follow up analysis.

### Multiplex test against SARS-CoV-2 epitopes

Serological status was evaluated using a proprietary, fluorescent bead-based multiplex assay, developed by the Cornell Animal Health Diagnostic Center, Serology Laboratory. The assay was developed and validated for detection of human antibodies (32) and was modified for antibody measurement in cats by using an anti-cat detection reagent. The feline assay evaluates antibody reactivity against three SARS-CoV-2 domains (NP, RBD, and S1). Based on validation with human serum, reactivity against NP or RBD is considered positive, while reactivity against S1 is considered cross-reactivity and not specific for previous SARS-CoV-2 exposure.

### Viral neutralization

Vero E6 cells, obtained originally from the American Type Culture Collection (ATCC) were seeded 24-hours prior to infection, under BSL-2 conditions, at a density of 2×10^4^ cells/well on a flat-bottom 96-well plate in 100 µL of Dulbecco’s modified Eagle medium (DMEM) (Cellgro) supplemented with 25 mM HEPES (Cellgro) and 10% HyClone Fetal Clone II (GE) at 37 °C and 5% CO_2_. For neutralization assays, under BSL-3 conditions, each heat inactivated sample was 4-fold serially diluted in Eagle’s Minimum Essential Medium (EMEM) (Cellgro) supplemented with 4% heat inactivated fetal bovine serum (FBS) (Gibco) and mixed with an equal volume of SARS-CoV-2 isolate USA-WA1/2020 from the Biological and Emerging Infections Resources Program (BEI Resources) for a final virus concentration of 100 TCID_50_. Dilutions were performed in triplicate. Following a one-hour incubation at 37 °C, media was removed from the Vero E6 cells and replaced with the serum/virus mixture. Plates were incubated for 72 hours at 37 °C and 5% CO_2_. After incubation, serum/virus was removed, and plates were fixed with 4% paraformaldehyde for 30 minutes and then stained with crystal violet for 10 minutes. The plates were washed with water and then allowed to dry. The neutralization titer was determined as the highest dilution in which at two or three of the wells were not-infected, based on visualization of the crystal violet staining. Samples were considered negative if there was no obvious viral neutralization at the 1/4 dilution.

### FCoV Serology

FCoV serology was performed by the Animal Health Diagnostic Center at Cornell University, utilizing a commercial ELISA for antibody detection, reported as a signal to noise (S/N) ratio and then classified as positive, negative, or inconclusive.

### Serology data mapping

Each sample had a metadata indicating geolocation and individual geolocation was analyzed and mapped using the Free and Open Source QGIS 3.34.12 (Free Software Foundation, Inc.).

### Statistical analysis

Unpaired student’s T tests and Fisher exact tests were performed to determine statistical significance between seropositivity of age and sex in GraphPad Prism 9 (San Diego, CA). A P-value of 0.05 was considered significant.

## Results

### Seropositivity

Out of the 79 samples collected, 13 cats (16.46%), tested positive for SARS-CoV-2 (Table 1, Figure 1). To further confirm these positive cases, we conducted live virus neutralization tests. Of the thirtheen seropositive samples, seven cats (S115, S123, S117, S149, S151, S166, and S167) exhibited neutralizing antibodies with titers ranging from 1/16 to 1/4096 (Table 1). Cats S115 and S123, sampled on 7/11/2020 and 7/18/2020 respectively, had shown respiratory signs in early March 2020. These two, exhibited the highest viral neutralization test (VNT) titers of 1/1024 and 1/4096, and were co-housed and owned by an individual who tested positive by serology for SARS-CoV-2 in April 2020. Cats S166 and S167, also which shared owner and living environment, were part of a household where other human housemates tested positive for SARS-CoV-2. Cats S149 and S151 were co-housed, but there was no reported information regarding their potential exposure to SARS-CoV-2. All these cats were kept indoors, except for S117, which was partially outdoor and inhabited in a community grocery store (Table 2). As for the remaining 6 seropositive cats (S113, S152, S153, S155, S157, and S158), although seropositive in the multiplex test, they tested negative in the VNT. Cat S153 had a travel history to a medical facility between May 5^th^ and 7^th^ 2020, and considering the community spread of SARS-CoV-2 in New York City at this time, the cat may have been exposed during its stay at the medical facility (Figure 1). In addition, we also performed viral neutralization with the 19 seronegative samples via multiplex. All the 19 seronegative cats were also negative by viral neutralization test.

**Table 1.** Multiplex ELISA and virus neutralization test (VNT)

| cat ID | NP | RBD | S1 | multiplex result | VNT result |
| --- | --- | --- | --- | --- | --- |
| S167 | 1889 | 8684 | 859 | positive | 1/16 |
| S115 | 6294 | 17400 | 967 | positive | 1/1024 |
| S117 | 714 | 5294 | 688 | positive | 1/256 |
| S149 | 7247 | 10862 | 1828 | positive | 1/256 |
| S151 | 8959 | 10139 | 1735 | positive | 1/256 |
| S123 | 9040 | 13408 | 1971 | positive | 1/4096 |
| S166 | 7055 | 3208 | 628 | positive | 1/64 |
| S101 | 487 | 1585 | 1759 | crossreactive (neg) | Neg |
| S103 | 719 | 1455 | 1616 | crossreactive (neg) | Neg |
| S105 | 1709 | 1583 | 1786 | crossreactive (neg) | Neg |
| S108 | 3544 | 2794 | 3290 | crossreactive (neg) | Neg |
| S109 | 4487 | 6151 | 7401 | crossreactive (neg) | Neg |
| S113 | 7044 | 1787 | 1783 | positive | Neg |
| S114 | 2731 | 699 | 665 | negative | Neg |
| S116 | 416 | 534 | 483 | negative | Neg |
| S127 | 892 | 766 | 762 | negative | Neg |
| S131 | 568 | 559 | 586 | negative | Neg |
| S132 | 453 | 538 | 576 | negative | Neg |
| S135 | 2207 | 754 | 743 | negative | Neg |
| S141 | 2391 | 4861 | 5135 | crossreactive (neg) | Neg |
| S145 | 3603 | 6386 | 6415 | crossreactive (neg) | Neg |
| S152 | 4572 | 2633 | 2820 | positive | Neg |
| S153 | 7295 | 2319 | 2255 | positive | Neg |
| S155 | 4806 | 415 | 478 | positive | Neg |
| S157 | 5332 | 718 | 982 | positive | Neg |
| S158 | 5201 | 901 | 1035 | positive | Neg |
| S159 | 2711 | 829 | 1000 | negative | Neg |
| S160 | 4513 | 1953 | 1970 | crossreactive (neg) | Neg |
| S163 | 4216 | 4241 | 4485 | crossreactive (neg) | Neg |
| S165 | 4228 | 887 | 1067 | crossreactive (neg) | Neg |
| S170 | 2860 | 1129 | 923 | negative | Neg |
| S172 | 3888 | 1377 | 1276 | crossreactive (neg) | Neg |
| S180 | 1897 | 1780 | 1909 | crossreactive (neg) | Neg |
| S181 | 2454 | 958 | 797 | negative | Neg |
| S102 | 695 | 938 | 1028 | negative |  |
| S104 | 501 | 1265 | 1299 | negative |  |
| S106 | 376 | 942 | 997 | negative |  |
| S107 | 469 | 1074 | 1040 | negative |  |
| S110 | 537 | 458 | 707 | negative |  |
| S111 | 568 | 521 | 808 | negative |  |
| S112 | 818 | 902 | 1288 | negative |  |
| S118 | 1260 | 911 | 955 | negative |  |
| S119 | 1451 | 1151 | 1252 | negative |  |
| S120 | 404 | 497 | 528 | negative |  |
| S121 | 543 | 590 | 518 | negative |  |
| S122 | 942 | 592 | 566 | negative |  |
| S124 | 389 | 527 | 546 | negative |  |
| S125 | 590 | 771 | 754 | negative |  |
| S126 | 727 | 744 | 778 | negative |  |
| S128 | 561 | 570 | 564 | negative |  |
| S129 | 728 | 428 | 490 | negative |  |
| S130 | 388 | 531 | 561 | negative |  |
| S133 | 411 | 503 | 471 | negative |  |
| S134 | 567 | 547 | 575 | negative |  |
| S136 | 712 | 800 | 936 | negative |  |
| S137 | 448 | 476 | 543 | negative |  |
| S138 | 886 | 865 | 982 | negative |  |
| S139 | 1060 | 758 | 850 | negative |  |
| S140 | 1197 | 745 | 836 | negative |  |
| S142 | 820 | 712 | 754 | negative |  |
| S143 | 594 | 731 | 819 | negative |  |
| S144 | 261 | 387 | 380 | negative |  |
| S146 | 989 | 664 | 629 | negative |  |
| S147 | 326 | 264 | 275 | negative |  |
| S148 | 540 | 822 | 747 | negative |  |
| S150 | 3037 | 873 | 736 | negative |  |
| S154 | 2369 | 304 | 307 | negative |  |
| S156 | 610 | 727 | 794 | negative |  |
| S169 | 1406 | 478 | 479 | negative |  |
| S162 | 1784 | 623 | 670 | negative |  |
| S161 | 3008 | 560 | 603 | negative |  |
| S164 | 2777 | 845 | 909 | negative |  |
| S168 | 1484 | 659 | 750 | negative |  |
| S171 | 1749 | 965 | 897 | negative |  |
| S173 | 2427 | 1277 | 1360 | negative |  |
| S174 | 1969 | 1135 | 1027 | negative |  |
| S177 | 1081 | 947 | 785 | negative |  |
| S178 | 2044 | 1018 | 895 | negative |  |
| S179 | 1303 | 1059 | 859 | negative |  |

**Table 2.**
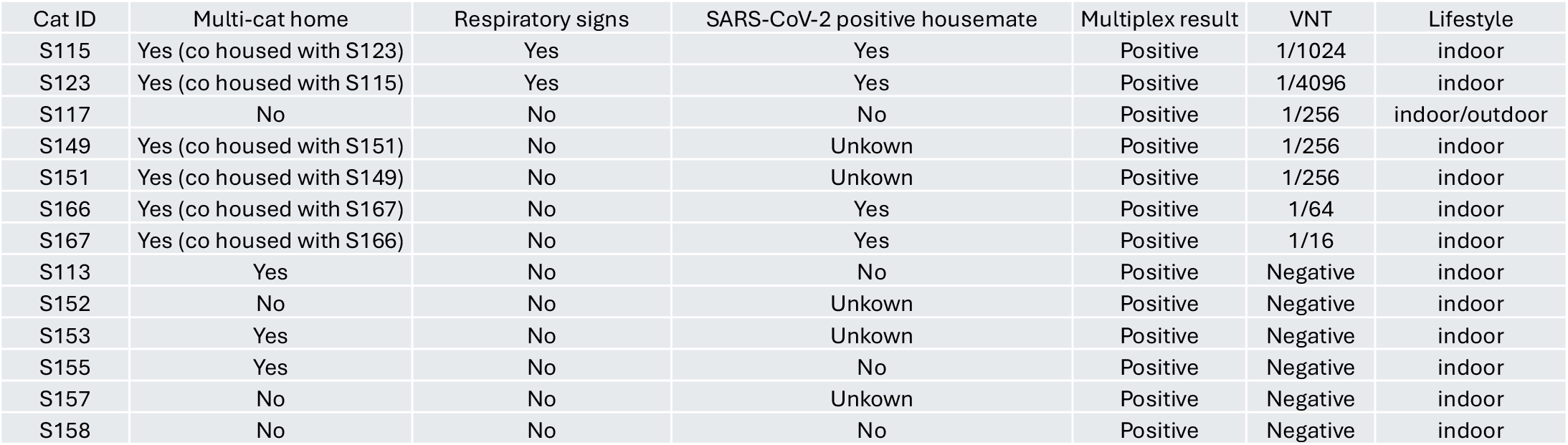
Association of seropositivity with respiratory signs.

| Cat ID | Multi-cat home | Respiratory signs | SARS-CoV-2 positive housemate | Multiplex result | VNT | Lifestyle |
| --- | --- | --- | --- | --- | --- | --- |
| S115 | Yes (co housed with S123) | Yes | Yes | Positive | 1/1024 | indoor |
| S123 | Yes (co housed with S115) | Yes | Yes | Positive | 1/4096 | indoor |
| S117 | No | No | No | Positive | 1/256 | indoor/outdoor |
| S149 | Yes (co housed with S151) | No | Unkown | Positive | 1/256 | indoor |
| S151 | Yes (co housed with S149) | No | Unkown | Positive | 1/256 | indoor |
| S166 | Yes (co housed with S167) | No | Yes | Positive | 1/64 | indoor |
| S167 | Yes (co housed with S166) | No | Yes | Positive | 1/16 | indoor |
| S113 | Yes | No | No | Positive | Negative | indoor |
| S152 | No | No | Unkown | Positive | Negative | indoor |
| S153 | Yes | No | Unkown | Positive | Negative | indoor |
| S155 | Yes | No | No | Positive | Negative | indoor |
| S157 | No | No | Unkown | Positive | Negative | indoor |
| S158 | No | No | No | Positive | Negative | indoor |

**Figure 1.**
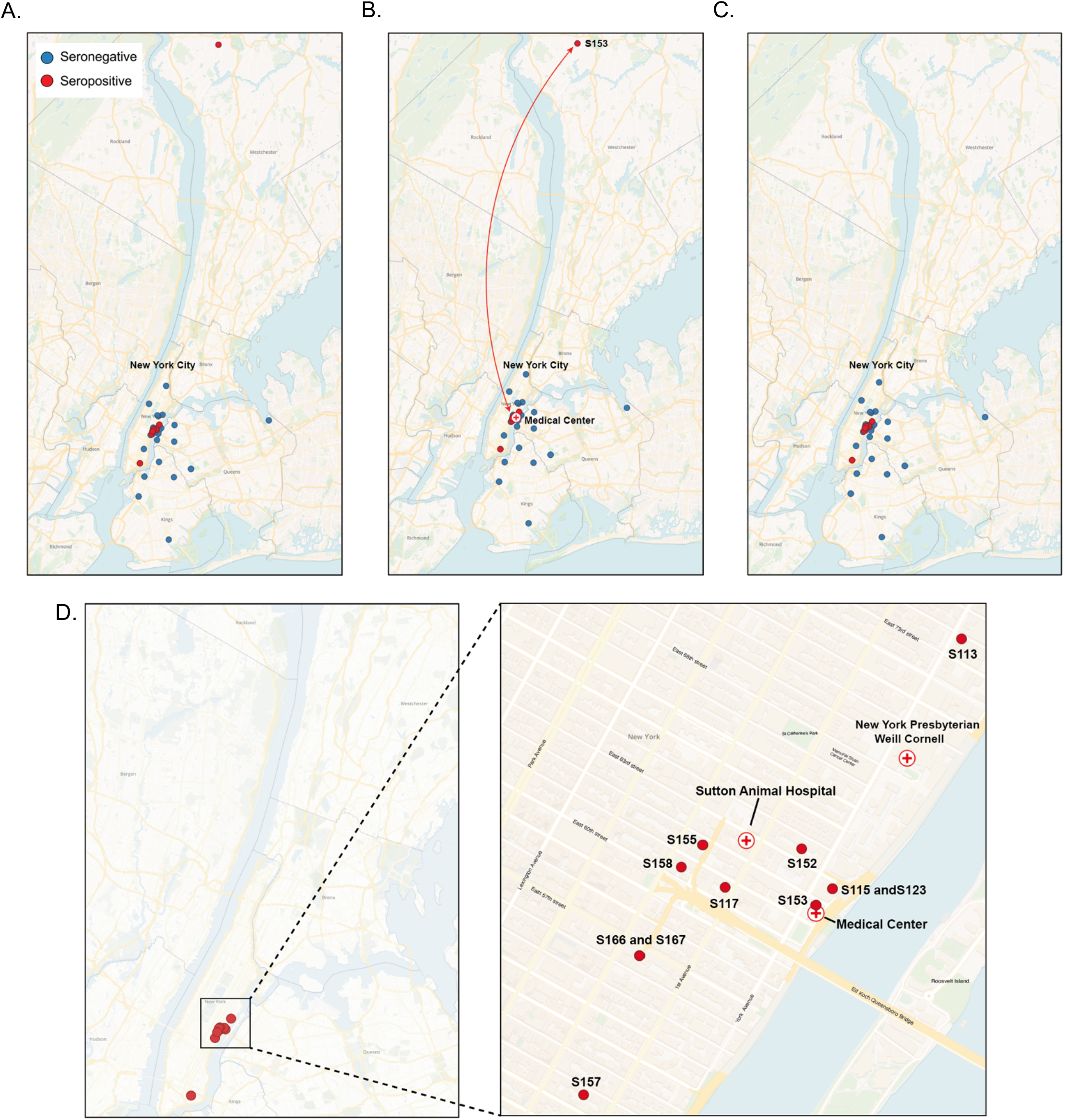
Location of sampled cats. (A) Location of all sampled cats. Blue circles corresponds to seronegative cats and red circles corresponds to seropositive cats. (B) Depitction of the travel history of cat S153. (C) Location of all sampled cats including travel history. (D) Enlarged view of location of seropositive cats.

### Association with sex, age, and breed

There was no observed association between cat age and sex with seropositivity in our analysis. Specifically, 84.8% of female cats tested negative, while 15.2% were positive. Among male cats, 81.8% were negative and 18.2% tested positive. Most of the cats we sampled were domestic long/short hair, therefore, most of the seropositive cats were domestic long/short hair (Figure 2).

**Figure 2.**
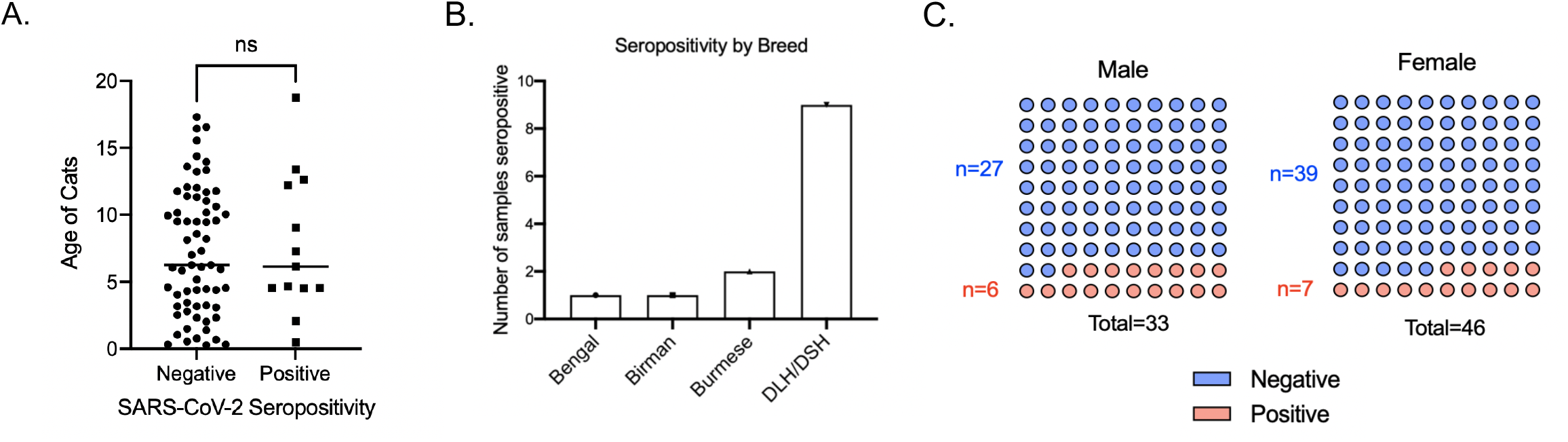
Association of seropositivity with age, breed, and sex. (A) Association between age of cats and seropositivity of cats. Each individual dot indicates individual cats. Statistical analysis was performed using unpaired student’s T test. (B) Seropositivity of different breeds of cats sampled. DLH: Domestic long hair. DSH: Domestic short hair. (C) Number of male and female cats sampled and their seropositivity. Statistical analysis was performed using Fisher exact test.

### Association with time of sample collection

The collection of samples was conducted over different months, and the seropositivity rates varied accordingly. The highest seroprevalence was observed during the fall of 2020, with 7 out of 16 cats testing positive (43.75%). In the summer of 2020, 4 out of 43 (9.3%) cats tested positive. During the winter of 2021, 2 out of 9 cats tested positive (22.2%). However, all samples collected during the spring of 2021 and summer of 2021 yielded negative results. (Figure 3).

**Figure 3.**
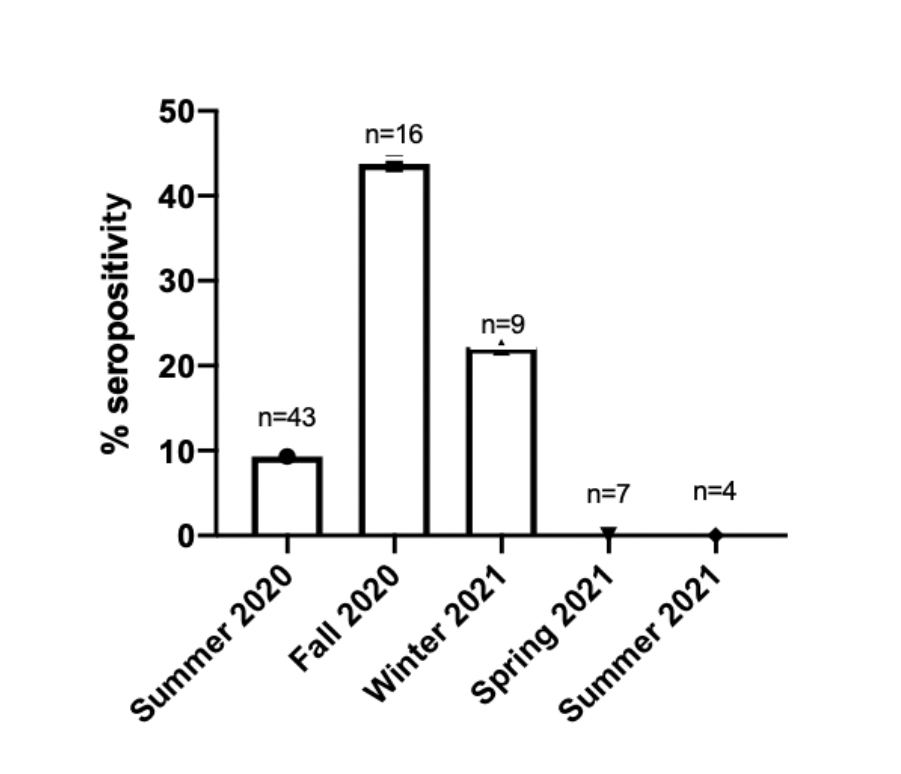
Association of seropositivity with time of sample collection. Number of seropositive samples based on time of sample collection. Samples were collected seasonally as follows: Summer (June, July, August), Fall (September, October, November), Winter (December, January, February), and Spring (March, April, May).

### Longitudinal samples

There were 12 longitudinal samples. Sera were collected from the same cat two to three times over a period. Three of these cats were serologically positive by multiplex ELISA and viral neutralization test. For all three samples, the viral neutralization titer decreased over time, but even a year after initial detection, it maintained a relatively high level (Table 3).

**Table 3.** Longitudinal samples.

| Repeat samples | Multiplex result | VNT result | Date collected |
| --- | --- | --- | --- |
| S117 | positive | 1/256 | 7/21/20 |
| S117 Repeat | low positive | 1/16 | 7/16/21 |
| S123 | positive | 1/4096 | 7/18/20 |
| S123 Repeat 1 | positive | 1/256 | 1/9/21 |
| S123 Repeat 2 | positive | 1/256 | 7/17/21 |
| S115 | positive | 1/1024 | 7/11/20 |
| S115 Repeat | positive | 1/256 | 1/9/21 |
| S121 | negative |  | 7/17/20 |
| S121 Repeat | low positive | negative | 1/9/21 |
| S142 | negative |  | 8/22/20 |
| S142 Repeat | low positive | negative | 1/5/21 |
| S153 | positive | negative | 9/5/20 |
| S153 Repeat | negative |  | 1/9/21 |
| S131 | negative | negative | 8/11/20 |
| S131 Repeat | negative |  | 2/9/21 |
| S107 | negative |  | 6/14/20 |
| S107 Repeat | negative |  | 2/27/21 |
| S102 | negative |  | 6/14/20 |
| S102 Repeat | negative |  | 3/13/21 |
| S134 | negative |  | 8/25/20 |
| S134 Repeat | negative |  | 7/24/21 |
| S135 | negative | negative | 8/25/20 |
| S135 Repeat | negative |  | 7/24/21 |
| S137 | negative |  | 8/25/20 |
| S137 Repeat | negative |  | 7/24/21 |

### FCoV serology

Among the 67 samples tested for FCoV, 34 (50.74%) tested positive, 20 (29.85%) negative, and 13 (19.4%) were inconclusive. The high prevalence of FCoV among these samples can be due to the fact that most of the cats were adopted from breeders or shelters, typically environments with multiple cats (Figure 4A). Among the 13 cats that tested positive for SARS-CoV-2, six were also found to be positive for FCoV (Figure 4B).

**Figure 4.**
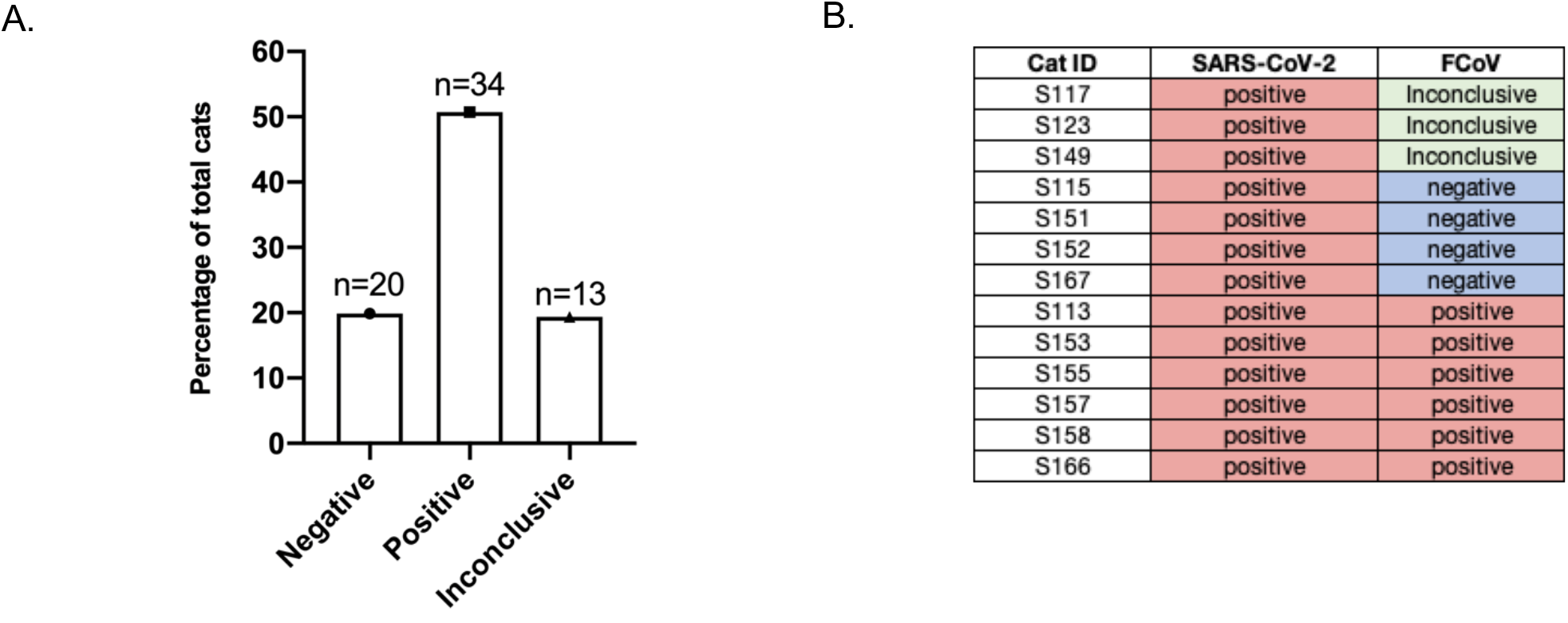
Prevalence of FCoV in feline sera collected. (A) Number of cats that tested negative, positive, and inconclusive for FCoV. (B) Table of SARS-CoV-2 seropositive cats and corresponding FCoV serology results.

## Discussion

In this study, we investigated the seroprevalence of SARS-CoV-2 in domestic cats during the early stages of the COVID-19 pandemic, highlighting the impact of the virus on this specific animal population. Although this study did not address aspects of viral shedding, our results suggest that cats were highly susceptible to SARS-CoV-2 infection during the early pandemic, and were capable of seroconverting and developing a sustained antibody response against SARS-CoV-2 upon infection.

Using a highly sensitive and specific fluorescent bead-based multiplex assay, we found an overall prevalence of 16.46% (13/79) seropositive animals for the study period, with a prevalence of 44% in the Fall of 2020. In comparison to other serosurveillance studies conducted on cats in the United States during the early phase of the pandemic (25–29), which range from 0.4% - 14% seropositivity, our study reports the highest seropositivity to date, which may be due to the geographic location and timing of our sampling—conducted while New York City was the epicenter of the COVID-19 pandemic in the United States.

The 6/13 cats tested positive on the fluorescent bead-based multiplex assay but negative on the VNT. From these, cats S155 and S158 exhibited decreased appetites. Cat S153 was admitted to a veterinary hospital in May 2020, given the timing of the stay, this travel history may have resulted in exposure to circulating SARS-CoV-2. The observed discrepancy between the multiplex assay and VNT results could likely be a result of the different analytical sensitivities of the two assays with lower limits of detection in the low pg/mL versus high ng/mL ranges for multiplex and VNT assays, respectively. The inconsistent results could also originate from either a false positive in the multiplex or a false negative in the viral neutralization test, or that the antibodies present were not neutralizing antibodies and thus only detected by the multiplex assay.

One notable finding from this study was the remarkably high seroprevalence of SARS-CoV-2 in cats during the fall of 2020, which reached 43.75% of the surveyed cats, aligning with the surge of human COVID-19 cases in NYC. This occurred in mostly indoor cats and at a time when social contact of their owners in the general population was heavily restricted. While we did not collect any information on the occupation of the owners or their health status, we note that the sampling site (Sutton Animal Hospital) is located in close proximity to a major medical complex (New York Presbyterian/Weill Cornell Medicine) that handled large numbers of human COVID-19 patients during the early phase of the pandemic; this proximity suggests a high level of asymptomatic spread in this restricted human population, and subsequently to their pets. Additionally, our samples were collected in the context of a “lock-down” that was imposed in New York City, at least during the early phase of the sampling period; the exception to more extended human contact was cat S117, who inhabited a community grocery store. Cats remain an interesting comparative model to study both serological and genomic/evolutionary responses to SARS-CoV-2 (33). As most cats were housed individually, our data are limited with respect to understanding cat-to-cat transmission.

In our study, we observed a high prevalence (50.74%) of feline coronavirus (FCoV) in owned domestic cats. FCoV is highly prevalent in feline populations especially in multi cat environments(34–41). Highly pathogenic variants of FCoV can cause feline infectious peritonitis (FIP), which is typically lethal in cats unless treated with antiviral drugs (5,42,43). Cats sampled in this study were predominantly indoor pets, often living alone with their owner or with just one other cat. Since the majority of cats sampled were from single-cat households and were indoor cats, they were likely exposed to FCoV prior to adoption from shelters or purchase from breeders, where they were in environments with multiple cats. Among the 13 cats seropositive for SARS-CoV-2, six also tested positive for FCoV. Considering the early pandemic timing of our sample collection, we infer that these cats were exposed to FCoV before encountering SARS-CoV-2. There is an over-arching question whether pre-exposure to other coronaviruses protects against SARS-CoV-2. The limited sample size in our study does not allow statistical significance to be reached, underscoring the need for additional research to validate this observation. The likelihood of cross-reactivity between FCoV and SARS-CoV-2 in serological tests is minimal, as prior studies have demonstrated an absence of cross-reactivity in FCoV antibodies against SARS-CoV-2 (44–46).

As with all studies of SARS-CoV-2, it is important to assess the individual virus lineages (24), especially with the Omicron variant, as substantial virological changes have occurred as the pandemic has proceeded. Based on the timing of the study, we sampled infections caused predominantly by pre-VOC SARS-CoV-2 (from the B.1 lineage, spanning both S:D614G and S:614D). In line with experimental studies of both “early” and “late” SARS-CoV-2 isolates, our study shows that cats can be readily infected with SARS-CoV-2 in natural populations, even where these populations are indoor cats, housed individually or in small groups.

The capacity of SARS-CoV-2 to infect a broad range of animal species has heightened concerns about reverse zoonotic transmission. The expansion of this viral reservoir could lead to an accumulation of mutations, potentially resulting in spillback events to the human population. Such events could give rise to variants that are either more transmissible or resistant to existing vaccines. Documented cases of reverse zoonotic transmission from animals to humans have been observed in minks, deer, and domestic cats (9,14,47). While our data did not address and understanding cat-to-human transmission. study underscores the critical need for a comprehensive cross-species approach to virus transmission, which includes rigorous and consistent surveillance measures across companion animals, livestock, and wildlife populations.

In this study, we did not directly collect information of owner exposure for SARS-CoV-2. As a result, we have incomplete data regarding the potential exposure of cats to SARS-CoV-2. It is plausible that some seronegative cats were exposed to SARS-CoV-2 but did not get infected or develop detectable antibodies. Therefore, the extent to which cats exposed to SARS-CoV-2 naturally gets infected or seroconvert remains unclear in our study.

In conclusion, our study underscores the importance of considering companion animals, particularly cats, in the broader epidemiological landscape of SARS-CoV-2. It highlights the necessity of ongoing research to understand the dynamics of interspecies transmission and the implications for both animal and human health. This research contributes to the growing body of evidence advocating for a One Health approach in managing and understanding the complexities of the COVID-19 pandemic.

## Acknowledgements

Funding for this study was provided by the Cornell University Office of the Vice-Provost for Research. AES was supported by the National Institutes of Health Comparative Medicine Training Program T32OD011000. AC was supported by grant T32EB023860 from the National Institute of Biomedical and Bioengineering.

This study was approved by The Cornell University Institutional Review Board for Human Participant Research (IRB) Office (IRB0009690; Serological study of coronavirus in cats) as an exempt protocol.

We thank Ana Bento and the members of the Whittaker Lab, past and present, for their helpful discussions during the course of this work. We also thank Lauren Coyle from Sutton Animal Hospital for handling and sending the cat sera.

